# Calcium activation of cortical neurons by continuous electrical stimulation: Frequency-dependence, temporal fidelity and activation density

**DOI:** 10.1101/338525

**Authors:** Nicholas J. Michelson, James R. Eles, Alberto L. Vazquez, Kip A Ludwig, Takashi DY Kozai

## Abstract

Electrical stimulation of the brain has become a mainstay of fundamental neuroscience research and an increasingly prevalent clinical therapy. Despite decades of use in basic neuroscience research and the growing prevalence of neuromodulation therapies, gaps in knowledge regarding activation or inactivation of neural elements over time have limited its ability to adequately interpret evoked downstream responses or fine-tune stimulation parameters to focus on desired responses. In this work, *in vivo* two-photon microscopy was used to image neuronal calcium activity in layer 2/3 neurons of somatosensory cortex (S1) in male C57BL/6J-Tg(Thy1-GCaMP6s)GP4.3Dkim/J mice during 30 s of continuous electrical stimulation at varying frequencies. We show frequency-dependent differences in spatial and temporal somatic responses during continuous stimulation. Our results elucidate conflicting results from prior studies reporting either dense spherical activation of somas biased towards those near the electrode, or sparse activation of somas at a distance via axons near the electrode. These findings indicate that the neural element specific temporal response local to the stimulating electrode changes as a function of applied charge density and frequency. These temporal responses need to be considered to properly interpret downstream circuit responses or determining mechanisms of action in basic science experiments or clinical therapeutic applications.

**Significance Statement:** Microstimulation of small populations of neurons has the potential to ameliorate symptoms associated with several neurological disorders. However, the specific mechanisms by which microstimulation elicits therapeutic responses are unclear. This work examines the effects of continuous microstimulation on the local population of neurons surrounding the implanted electrode. Stimulation was found to elicit spatiotemporal neuronal responses in a frequency dependent manner. These findings suggest that stimulation frequency may be an important consideration for applications in research or therapy. Further research aimed at understanding these neuronal activation properties may provide insight into the mechanistic mode of action of continuous microstimulation.

## Introduction

Understanding the complexity of the nervous systems requires the ability to readout detailed measurements from controlled inputs (Adrian, 1928; Adrian & Bronk, 1928; Adrian & Matthews, 1928; Sherrington, 1910). While many new techniques and technologies are emerging, electrical stimulation remains one of the oldest and most widely used methods for directly interfacing and driving the nervous system(Du Bois-Reymond, 1884; Fritsch & Hitzig, 1870; Galvani, 1791; Galvani & Aldini, 1792; Volta, 1800). The ability to directly apply electrical potentials to individual nerves has helped identify biophysical properties of nerves and neurons(Eccles, 1953; Hodgkin & Huxley, 1952), and has been crucial in understanding the organization of the brain (Penfield & Boldrey, 1937; Spiegel, Wycis, Marks, & Lee, 1947). These studies also opened the door to electrical stimulation approaches for therapeutic applications (Delgado, Hamlin, & Chapman, 1952). Electrical stimulation has been used for hearing(Djourno, Eyries, & Vallancien, 1957; Doyle Jr, Doyle, Turnbull, Abbey, & House, 1963; L. R. House, 1987; W. F. House, 1976), vision (W. Dobelle & Mladejovsky, 1974; W. H. Dobelle, Mladejovsky, & Girvin, 1974), and somatosensory restoration (Cushing, 1909; Flesher et al., 2016) as well as for treating movement disorders (Alberts et al., 1965; Hassler, Riechert, Mundinger, Umbach, & Ganglberger, 1960; Sem-Jacobsen, 1966; Sem-Jacobsen, 1965), chronic pain (Gol, 1967; Mayer & Liebeskind, 1974; Richardson & Akil, 1977a, 1977b), and epilepsy (Marcos et al., 2000; Robert et al., 2010). More recently there has been an expansion of neuromodulation clinical trials, with over 1000 clinical trials registered on clinicaltrials.gov utilizing implanted or non-invasive electrodes to activate or inactivate nervous tissue. Some of these application include treatment of diverse conditions such as Tourette’s syndrome, obsessive compulsive disorder, depression, Alzheimer’s disease, and stroke. These clinical applications further motivate the development of improved chronically implanted electrical stimulation devices (LifeScienceAlley, 2015).

Despite its growing prevalence in scientific and clinical research, there is limited understanding of how electrical stimulation directly interacts with the complex milieu of neuronal and non-neuronal cells in the brain near the electrode. In acute mapping experiments of peripheral nerve fibers, short pulses of electrical stimulation were hypothesized to activate a small sphere of fibers in the local vicinity of the electrode (Ranck, 1975; Rattay, 1999; Stoney Jr, Thompson, & Asanuma, 1968; Tehovnik, 1996), with increases in amplitude increasing the volume of the sphere activated. When applying cathodic leading stimulation waveforms, increasing the amplitude could also cause a small zone of inactivation near the electrode tip, due to the generation of virtual anodes more distant from the tip, ultimately hyperpolarizing axons and preventing the propagation of any action potentials (Ranck, 1975). In contrast, recent experiments utilizing 2-photon calcium imaging to visualize the activation of neural elements in cortex near the electrode site found that electrical stimulation activated a sparse population of neuronal somas distant from the electrode site (Histed, Bonin, & Reid, 2009). These experiments demonstrated that electrical stimulation activates a much smaller volume of neural processes around the electrode tip, which in turn activate the connected soma (Histed et al., 2009).

Although changes in pulse amplitude are widely understood to alter the area of activation within the vicinity of the electrode, the extent to which different stimulation frequencies and pulse train durations affect local neuronal responses remains less explored. In this study, *in vivo* multiphoton calcium imaging of GCaMP6 transgenic mice was employed to elucidate how neuronal elements near an implanted electrode respond to different electrical stimulation frequencies over the course of a longer pulse train. Transgenic GCaMP6 mice allow for the probing of neuronal activity with high fidelity in a large proportion of cortical excitatory neurons without the need for exogenous calcium chelators. Stimulation was carried out at different frequencies using a 30 second burst of cathodic leading symmetric pulses, with a 50 μs pulse width/phase and pulse amplitudes within the established Shannon safety limits to avoid tissue damage (Shannon criteria of *k* between 0.12 and 1.3, 2.5 nC per phase). Based on the results of Histed et al (Histed et al., 2009) and Stoney et al (Stoney Jr et al., 1968), we hypothesized that both sparse and distributed activation would occur in cortex with continuous electrical stimulation at sufficient charge/phase. Our results support this hypothesis, but also demonstrate that different neural elements (neurites, soma) in the vicinity of the electrode differ in their responses as a function of time (stimulation onset, during, and after stimulation), space, and stimulation frequency. These data show a complex response profile near the electrode to electrical stimulation that may have significant implications for the design of electrical neuromodulation experiments and therapies. Hence, it is not surprising that different studies utilizing different charge/phase, frequencies, and time points yield conflicting results.

## Methods

### Surgery and electrode implantation

Transgenic mice C57BL/6J-Tg(Thy1 GCaMP6s)GP4.3Dkim/J (n=5, male, 22–28 g; IMSR Cat# JAX:024275, RRID:IMSR_JAX:024275) were used in this experiment. All surgical interventions were performed on adult mice (>4 weeks of age) that were housed in social housing with 12h light / 12h dark cycles and free access to food and water until acute experimentation. All animals were induced with an anesthetic mixture consisting of 75 mg/kg ketamine and 7 mg/kg xylazine administered intraperitoneally and updated with 40 mg/kg as needed. Rectangular craniotomies (~4 mm per side) were made over each somatosensory cortex and electrodes were implanted at a 30° angle as previously described (Eles et al., 2017; T. D. Kozai, Vazquez, Weaver, Kim, & Cui, 2012; Takashi D. Y. Kozai, Eles, Vazquez, & Cui, 2016; Takashi D Y Kozai, Jaquins-gerstl, Vazquez, Michael, & Cui, 2016; Michelson et al., 2018; S. M. Wellman & Kozai, 2018). Electrical stimulation was conducted through a single-shank 16-channel Michigan style functional silicon probes with 703 μm^2^ electrode sites (NeuroNexus Technologies, Ann Arbor, MI). In order to prevent mechanical strain-related Ca activity caused by microelectrode insertion (Eles, Kozai, Vazquez, & Cui, 2018) from influencing stimulation evoked GCaMP activity, the tissue was allowed to rest 20 minutes after insertion prior to stimulation experiments. All experimental protocols were approved by the University of Pittsburgh, Division of Laboratory Animal Resources and Institutional Animal Care and Use Committee in accordance with the standards for humane animal care as set by the Animal Welfare Act and the National Institutes of Health Guide for the Care and Use of Laboratory Animals.

### Two-photon Imaging

A two-photon laser scanning microscope (Bruker, Madison, WI) and an OPO laser (Insight DS+; Spectra-Physics, Menlo Park, CA) tuned to a wavelength of 920 nm at 15 mW was used for the entirety of this study. A16X 0.8 numerical aperture water immersion objective lens (Nikon Instruments, Melville, NY) was selected for its 3 mm working distance. Imaging was carried out over a 407 x 407 μm ROI, collected with a 512 x 512 pixel matrix. Before imaging sessions, animals were injected IP with 0.1 ml of 1mg/ml SR101 for visualization of blood vessels and electrode contacts. Imaging plane was positioned 10 μm above the electrode site, 120-150 μm below the surface of the brain. In order to image neural dynamic during electrical stimulation, time series images were recorded at 30 Hz and 2x optical zoom. The imaging plane for time series was set right over the electrode contact, so that the stimulating contact is visible during imaging session.

### Electrical stimulation

Concurrent stimulation and imaging typically commenced 20 minutes after electrode implantation to allow time for displaced tissue to settle. An A-M Systems 2100, single channel Isolated Pulse Stimulator (A-M Systems, Sequim, WA) or TDT IZ2 stimulator on an RZ5D system (Tucker-Davis Technologies, Alachua, FL) were used to apply current controlled cathodic leading biphasic symmetric (50 μs each phase) electrical microsimulation at 50μA and frequencies of 10, 30, 50, 75, 90, 130 and 250 Hz. However, in one animal, the impedance of the stimulated site was an order of magnitude lower than that listed by the manufacturer, suggesting minor insulation failure at the electrode/insulation interface, as noted in previously (Prasad et al., 2014). This has the net effect of slightly increasing the electrode interface area, requiring greater current to achieve the same effective charge density seen in other animals. In this animal, 100 μA stimulation current was necessary to drive GCaMP activity. Amplitude of stimulation was selected based on minimum current required to evoke activation during two photon live scanning. Charge/phase, charge density/phase and k values were calculated based on the Shannon’s equation (Shannon, 1992). The *k* value used during stimulation was between −0.12 and 1.3, well within safety limits to avoid tissue damage. After initial collection of resting state activity for 2min, concurrent imaging and stimulation trials were acquired for 2 min at each frequency, which included 30 s of pre-stimulation baseline, 30s of stimulation and 60s of post-stimulation baseline. An additional 3 minutes was required between each trial to store the data after data collection. The two-photon microscope was synchronized to the stimulator via transistor-transistor logic (TTL) and 30 s stimulation was carried out after a 30 s delay from the start of the time series. Stimulation was carried out in increasing frequency and were not repeated in the same animal. However, in two animals, 10 Hz stimulation was repeated at the end to determine if earlier microstimulation trials change the neuronal response properties for subsequent trials. The second 10 Hz stimulation data showed similar activation patterns to the first 10 Hz stimulation data.

### Data analysis

#### Identification of activated neurons

The acquired time series images were converted to a single tiff image file using ImageJ (NIH). The tiff image was then analyzed using a custom MATLAB script (MATLAB, RRID:SCR_001622). Non-overlapping elliptical regions of interest (ROIs) were manually drawn around neurons to fully encompass their somas, which were identified by their morphology as bright rings of fluorescence (Fig 1b,d). ROIs were obtained using the CROIEditor MATLAB script, written by Jonas Reber (https://www.mathworks.com/matlabcentral/fileexchange/31388-multi-roi-mask-editor-class). Fluorescence intensity time-courses for each neuron were generated by averaging all pixels within a cell’s ROI. Baseline fluorescence intensity (F_0_) was calculated as the average signal during the 30-second pre-stimulation interval, and each time-course was then converted to changes in fluorescence (ΔF/F_0_). To identify activated neurons, each neuron’s ΔF/F_0_ signal during stimulation was compared with their pre-stimulation ΔF/F_0_ signal. A fluorescence threshold of the mean pre-stimulation signal + three standard deviations (SD) was then established, such that neurons which showed ΔF/F0 changes that exceeded the threshold during stimulation were considered to be activated by the electrical stimulation. While it is possible that the threshold selection might not have identified all active cells, GCaMP6s is very sensitive to neuronal activity even for single action potentials (>30% ΔF/F reported) (Chen et al., 2013). Our criteria was robust for detection of low levels of activity even in the presence of small, unwanted signal changes due to motion or photomultiplier tube (PMT) dark current. The spatial locations and the activation properties of these neurons were then quantified for further analysis.

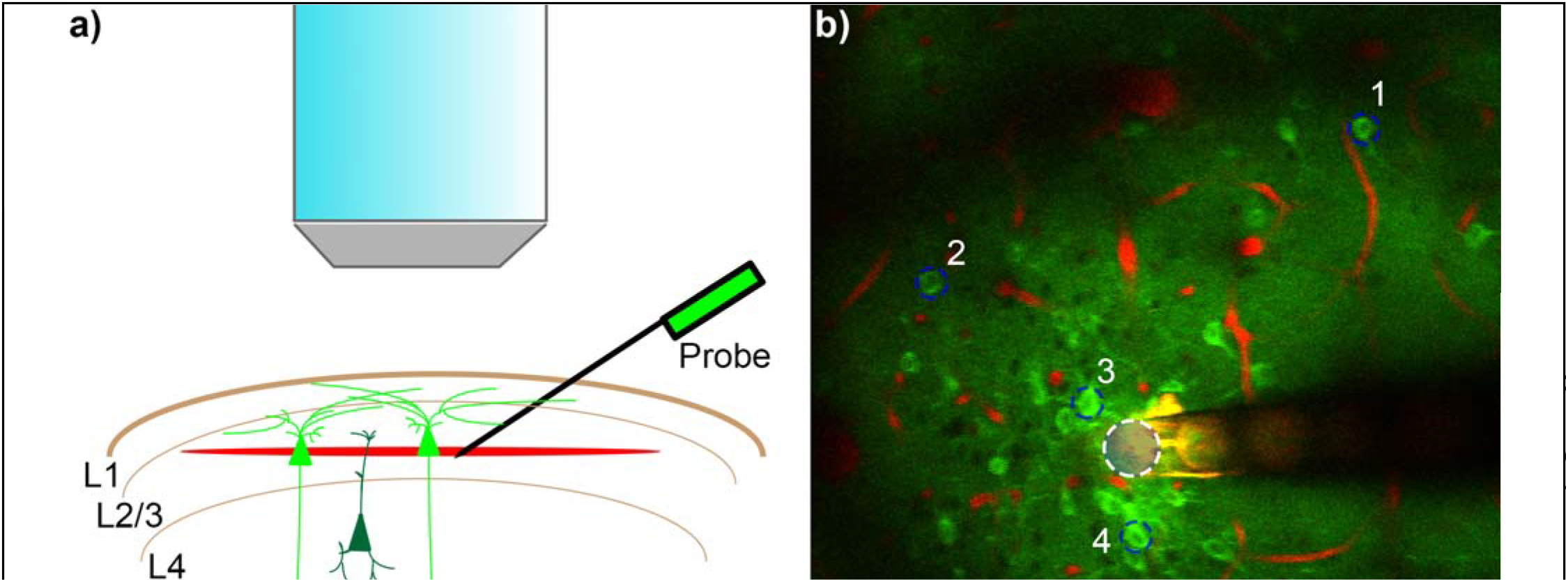

#### Neuropil Activation

Neuropil intensity was quantified by averaging the intensity level of a 10 x 25 μm region that did not contain neuronal somas. For each frequency, the neuropil was measured in the following distances from the edge of the electrode; 10-20, 30-40, 50-60, 100-110, 150-160, and 245-255 μm. Similarly, representative neuronal somas were selected at 15 and 115 μm from the electrode. F/F_0_ of GCaMP intensities were computed where F_0_ was the average GCaMP intensity of the 30 s pre-stimulus period.

#### Temporal activation pattern

Somas were classified as onset-responsive if they were activated during the first 0-2 seconds of the 30 second pulse train and ceased activation before the final 2 seconds of the pulse train (28-30 s). Somas were classified as steady-state-responsive neurons if they were active during 28-30 s post-stimulation onset. Duration of activity was calculated for each neuron by considering all time-points during stimulation which exceeded the intensity threshold. To examine the temporal characteristics of activated neurons in local vs distant or dense vs sparse groups, a weighted response time was estimated for each neuron. The ‘Activation Time’ was calculated as the time for which the cumulative ΔF/F sum reached 50% of the total ΔF/F sum over the 30s stimulation period. This expression describes the time it takes to cover half of the response energy. For example, a soma that reaches peak intensity at 0 seconds into the pulse train and maintains the exact same ΔF/F for all 30 s would have an ‘Activation Time’ of 15 s. Similarly, a soma that maintains a peak ΔF/F from 0 s to 5 s, but then falls to 20% of peak ΔF/F from 5 s to 30 s would have an ‘Activation Time’ of 5 s.

#### Spatial activation pattern

The spatial relationship between stimulation frequency and activity was determined by measuring the distance from the center of each active neuron to the nearest edge of the stimulating electrode site. The location of the center of the stimulating electrode site was visually identified from the same image series that was used to identify activated neurons and was confirmed with a high-resolution image taken immediately before the trial. The distances from the nearest edge of the stimulating electrode site to each activated neuron during the stimulation period were estimated for all animals and stimulation frequencies. The mean distance + 1SD was then calculated for each stimulation frequency and animal. Because the distance of onset and steady state activated neurons were considered for these calculations, distance thresholds remained fairly consistent across stimulation frequencies within animals. Therefore, the stimulation frequency with the minimum mean + 1SD for each animal was chosen to represent a distance threshold (139.7±18.8 μm for n=5), whereby cells in this region were considered local neurons, while those beyond the distance threshold were considered distant neurons.

#### Activation density

Density was examined as an independent measure by creating concentric circular bins, with an increasing radius of 20 μm drawn around the center of the electrode site. The number of fully encompassed, activated neurons within the area defined by each bin were counted from 0 to 30 s post stimulation onset, and then divided by the total area encompassed by that bin. Activated neurons were separated into dense or sparse populations, by establishing a threshold of mean density + 1 SD. To compare high and low-density activation patterns across stimulation frequencies, density thresholds were established for all stimulation frequencies, and the lowest mean density threshold was applied to all trials.

#### Statistical analysis

All data in boxplots show the sample median (eyes), 25^th^ and 75^th^ percentiles (top and bottom edges of box), 1.5 times the interquartile range (whiskers), and outliers (individual dots). To assess significance between stimulation frequencies, a repeated measures two-way ANOVA was performed (SPSS, RRID:SCR_002865), followed by post-hoc two-sample unequal variance t-test with a Bonferroni correction.

## Results

*In vivo* two-photon calcium imaging of ketamine/xylazine-anesthetized GCaMP6 transgenic mice was used to study the spatiotemporal activation pattern of neurons during microstimulation. Neurons in layer II/III of S1 somatosensory cortex were imaged as previously described (Michelson et al., 2018), and electrically stimulated continuously for 30 s at different frequencies (Fig 1a). GCaMPs are genetically encoded intracellular calcium indicators that largely reflect action potential firing (Chen et al., 2013; Dana et al., 2014; Greenberg, Houweling, & Kerr, 2008; Kerr & Denk, 2008; Sato, Gray, Mainen, & Svoboda, 2007; Smetters, Majewska, & Yuste, 1999). Because quiescent neurons have low GCaMP fluorescence, capillary visualization using 0.1 mg of sulforhodamin101 (SR101) was used to ensure the location of the electrode and visibility in the tissue with low laser power (< 20 mW), as previously described (Fig 1b) (Eles et al., 2017; T. D. Kozai et al., 2012; Takashi D. Y. Kozai et al., 2016; Takashi D Y Kozai et al., 2016; T. D. Y. Kozai et al., 2010; Michelson et al., 2018; S. M. Wellman & Kozai, 2018). Although GCaMP6 is more sensitive compared to other calcium indicators, such as Oregon green BAPTA, it has a longer decay constant that did not impair detection of activity(Chen et al., 2013). Evoked fluorescence traces show considerable variability in the intracellular calcium response to continuous stimulation (Fig 1c). Due to the binding constant, brightness, detector saturation, and temporal imaging resolution, it is not possible to quantify the calcium transients. The mouse line used in this study (GP4.3Dkim/J) has relatively high expression levels of GCaMP in layer II/III neurons (Dana et al., 2014), which was corroborated in a separate study, where this mouse line was imaged during a cortical spreading depression (Fig. 1d). The results demonstrate that sparse GCaMP activation noted during electrical stimulation is not due to sparse GCaMP expression in the genetic line.

The laminar probes were inserted with the electrode sites facing up and the imaging plane positioned approximately 10-15 μm above the electrode site in order to capture the neurons just above the stimulation site. The position of the imaging plane was confirmed through anatomical Z-stacks taken prior to the experiment. Co-registration of vasculature features outline of the implanted probe, and the electrode sites and traces were used to ensure that the imaging plane was just above the electrode site throughout the experiment. Electrical stimulation parameters matched previously employed protocols for neuroprosthetics and neuromodulation applications (De Balthasar et al., 2003; Kent & Grill, 2012; Kim et al., 2015; Shepherd & Javel, 1999; Wu, Levy, Ashby, Tasker, & Dostrovsky, 2001). Stimulation pulses were cathodic leading biphasic symmetric waveforms with pulse widths of 50 μs/phase (Lilly, Hughes, Alvord, & Galkin, 1955). Stimulation was performed for 30 s at a charge density and charge per phase below the Shannon safety limit for planar iridium electrode sites with 703 μm^2^ surface area (Shannon, 1992) (< 3nC/phase) (Cogan, Ludwig, Welle, & Takmakov, 2016; D. B. McCreery, Yuen, Agnew, & Bullara, 1994). In general, stimulation was delivered at 50 μA and 50 μs, which has been previously shown to be just above the sensitivity threshold (Kim et al., 2015). The stimulation frequency was varied over different trials (Fig. 2–3; 10, 30, 50, 75, 90, 130, 250 Hz). As expected from previous reports (Histed et al., 2009), electrical stimulation led to sparse, distributed activation patterns (Fig. 2). Frequencies of 50 Hz and 130 Hz were selected as they are the frequencies at which the Shannon/McCreery safety curve was developed and the average optimal therapeutic frequency for deep brain stimulation (DBS) in Parkinson’s Disease patients, respectively (Cogan et al., 2016; Shannon, 1992). These frequencies are compared to 10 Hz stimulation (Fig. 2).

**Figure 2:**
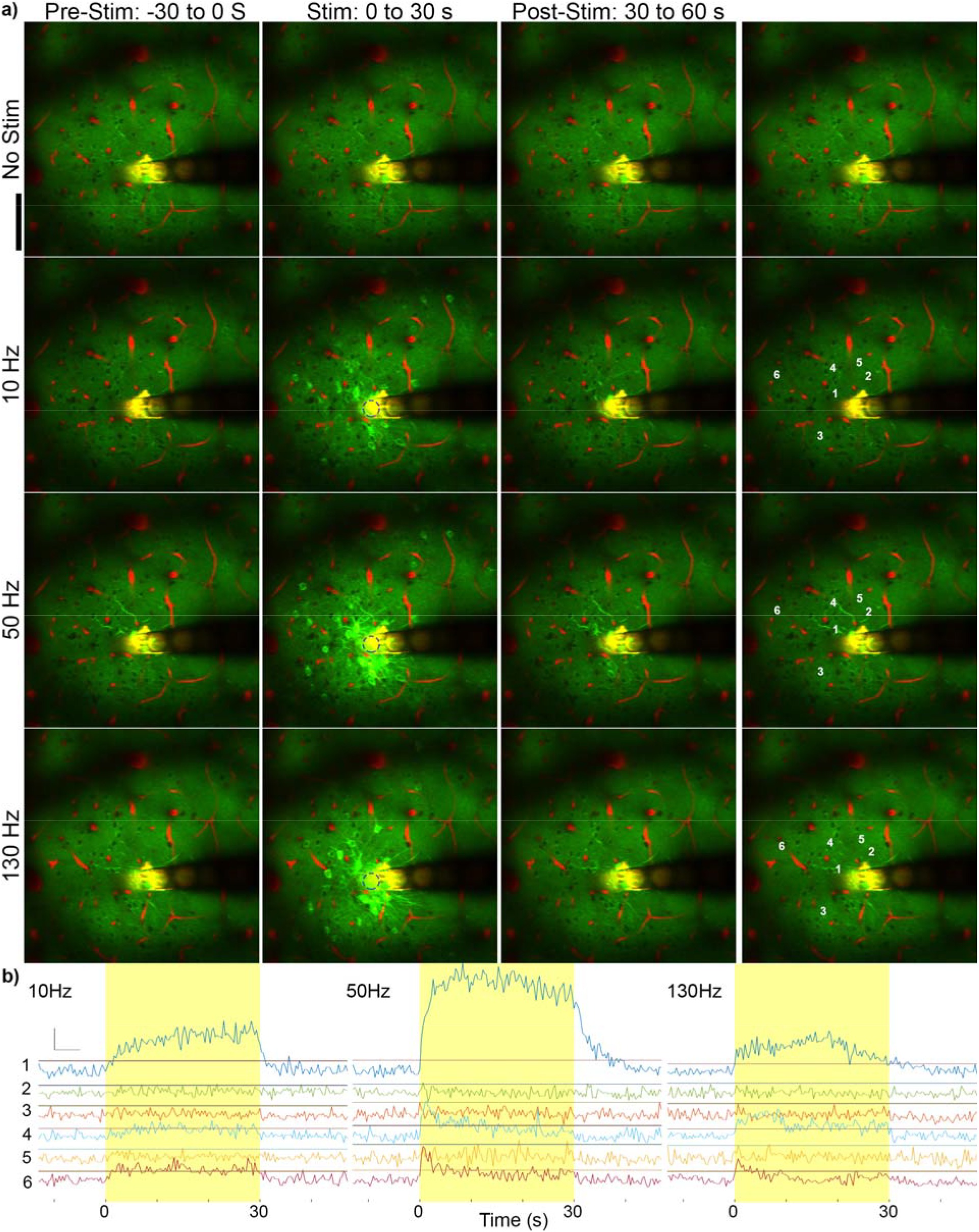
Stimulation frequency influences GCaMP activation pattern. a) Mean GCaMP imaging of 30 s prior to stimulation (left), during stimulation (middle), and post-stimulation (right) at no stimulation, 10 Hz, 50 Hz, and 130 Hz. Green indicates GCaMP labelled neurons and red indicates vasculature. The electrode sites and traces reflect fluorescence in both channels and therefore appear in yellow. Different patterns of activation can be observed at different frequencies, and GCaMP activity returns to baseline by 30 s poststimulus. Scale = 100 μm. k=-0.12. b) GCaMP activity of labeled cells with increasing distances from the electrode (by distance from the electrode edge: 1 = 15 μm; 2 = 33 μm; 3 = 51 μm; 4 = 54 μm; 5 = 64 μm; and 6 = 105 μm). Horizontal lines indicate threshold. Neurons 1, 4, and 6 show electrical stimulation induced GCaMP activity above threshold while 2, 3, and 5 do not.

**Figure 3:**
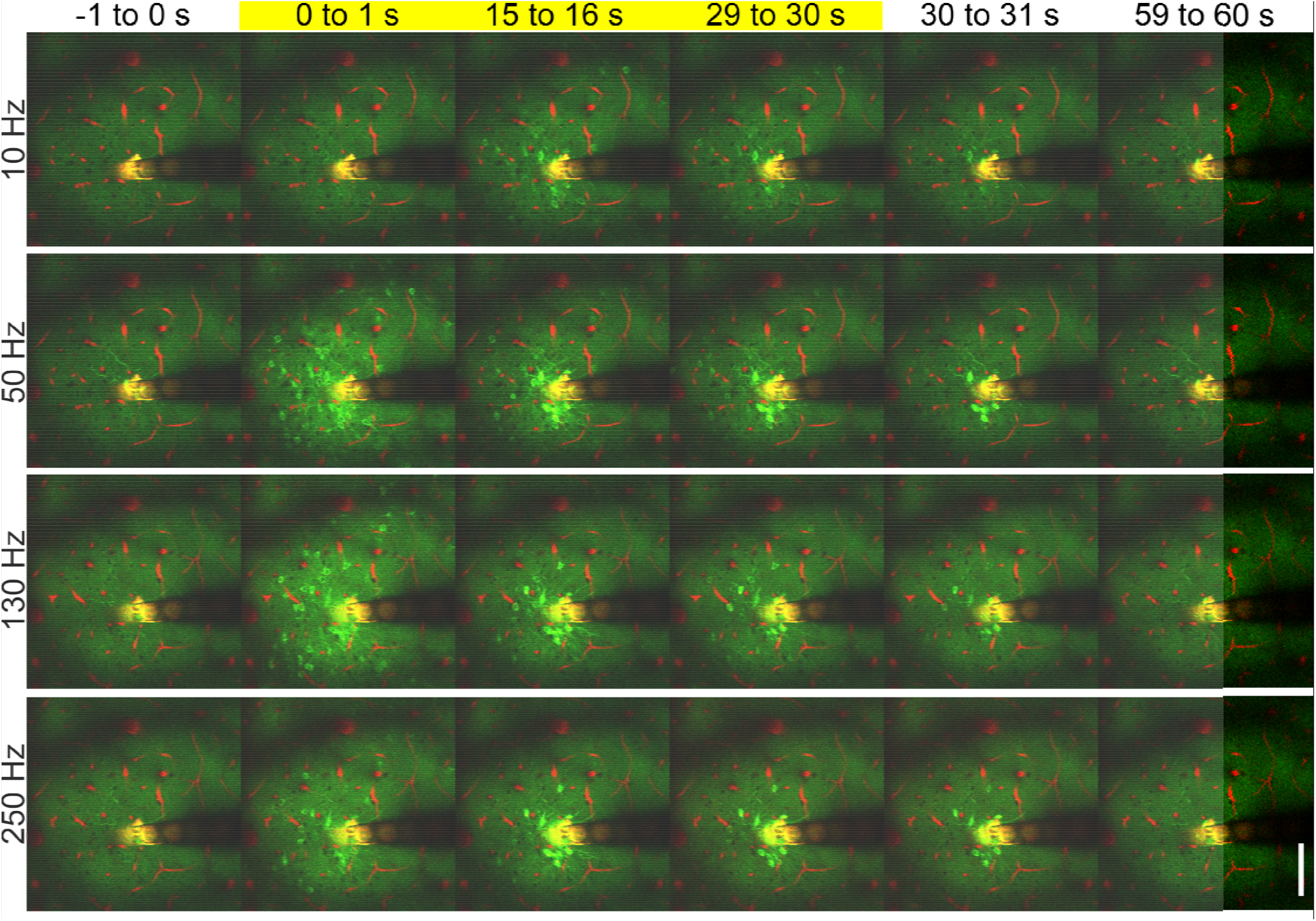
Continuous electrical stimulation evokes frequency dependent temporal activation. Mean GCaMP activity immediately before stimulation (−1 to 0s), at stimulation onset (0 to 1s), during stimulation (15 to 16s), immediately before stimulation termination (29 to 30s), immediately after termination (30 to 31s) and after stimulation (59 to 60 s). While the activation of GCaMP is lower at 10 Hz, GCaMP activation pattern is similar at onset across frequencies. However, the activation pattern at the end of the stimulation train (29 to 30 s) differs based on frequency of the stimulation. Images taken from the same animal and trials as figure 2. Scale = 100 μm. *k*=-0.12.

### Distinct GCaMP intensity falloff between neuronal somas and neuropil during stimulation

Neuropil activation as a function of distance from the electrode, time post-stimulation onset, and frequency of stimulation was also examined (Fig. 5). Since the neuropil is composed of neurites from surrounding GCaMP-expressing cells, it is understood to represent a spatial average of nearby responding and non-responding cells (Histed et al., 2009). In this analysis, regions of interest previously identified as isolated somas were removed, and the average changes in GCaMP intensity in the remaining volume were assessed. Across all frequencies tested, there appears to be a near linear fall off in stimulation evoked GCaMP intensity as a function of distance from the electrode (Fig. 5b red vs orange vs green lines). In contrast to the neuropil, example neurons near (15 μm) the electrode site show gradual increase in GCaMP intensity at 10 Hz and a rapid plateau between 30 to 90 Hz (Fig. 5b, blue; 15 μm). However, similar to the time course of distant soma activation (Fig. 5b, red; 115 μm), there is a peak in activation of the neuropil in the first five seconds post stimulation, with a decrease in activation thereafter. At 10 Hz the decrease in neuropil activation is slower across all distances, with the peak and subsequent decay being steeper (slope of F/F_0_) for higher frequencies. There is also a poststimulus residual activation for both the neuropil and local neurons (Fig. 5a b). This residual increase in activation can last up to 30 seconds or more post stimulation (Fig. 4b) for somas, but is typically shorter in duration for the neuropil (Fig. 5a).

**Figure 4:**
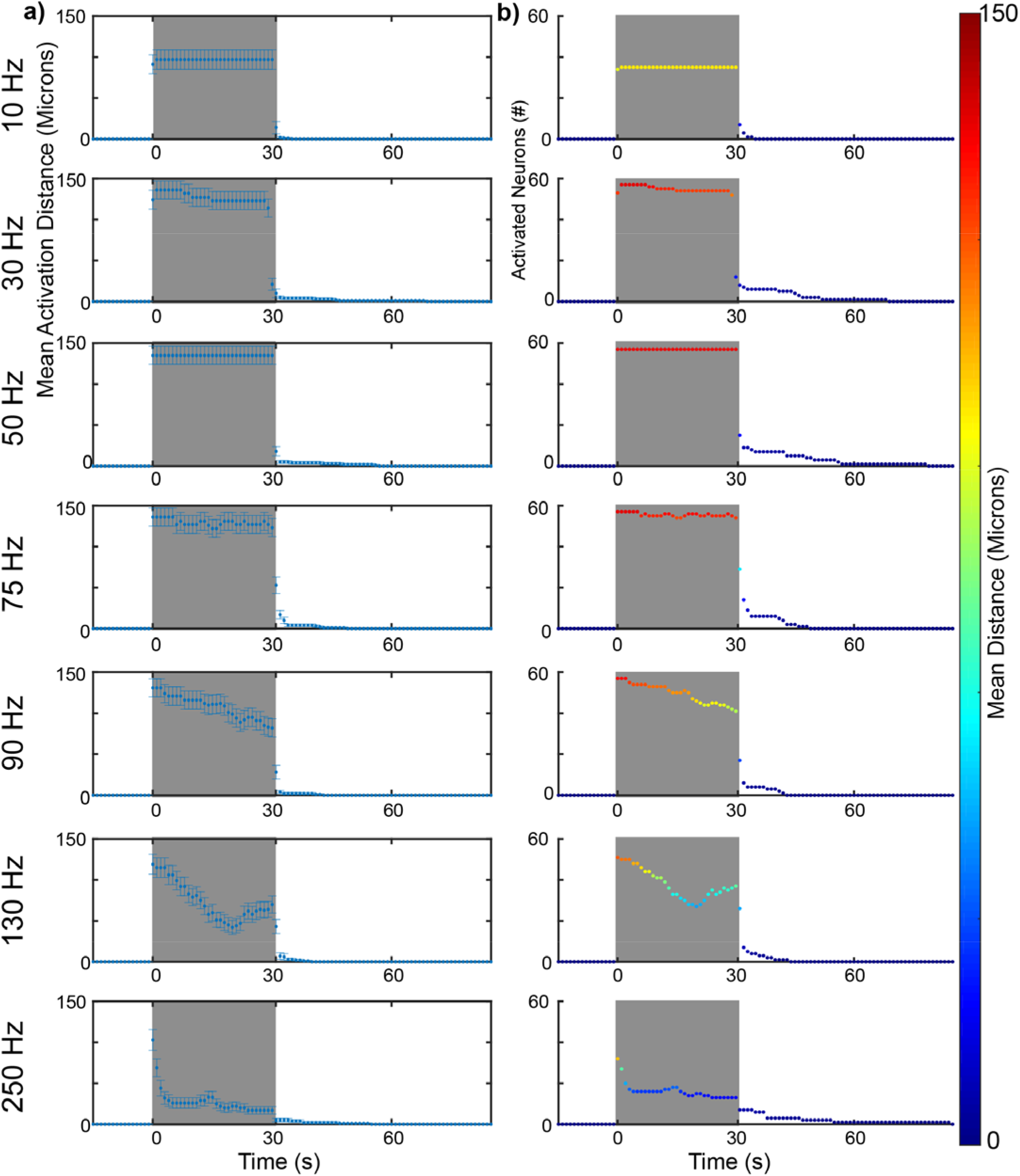
Spatial and temporal distribution of frequency dependent GCaMP activation in a representative animal. **a**) The average distance of all active neurons during electrical stimulation (grey box) decreases over time at higher frequencies. Error bars indicate standard error. **b**) The number of active neurons during electrical stimulation also decreases over time at higher frequencies. Mean activation distance from panel (a) is represented in color to show that electrical stimulation at higher frequencies (>90 Hz) leads to a decrease in the average distance of calcium activity over long stimulation pulse trains (color) and decrease in variance (a), suggesting the decrease in average distance is due to discontinued activity of distant neurons.

**Figure 5.**
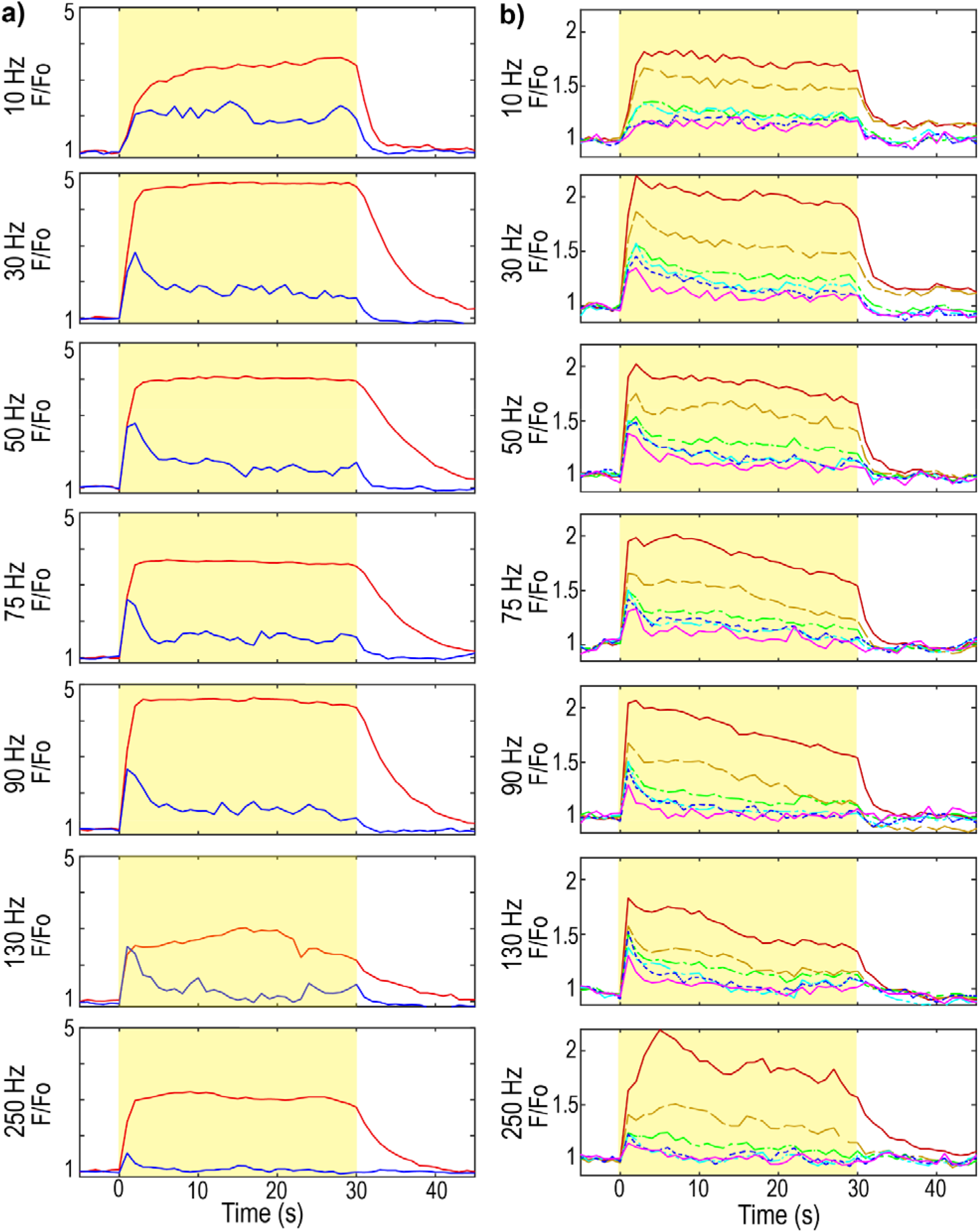
Neuronal soma activation and neuropil activation show frequency dependent temporal falloff. **a**) GCaMP activation in the soma at 15 μm (red) or 115 μm (blue) from the electrode site. Yellow indicates the stimulation. F_0_ is the mean intensity of the pre-stimulus period. Calcium activity was averaged into 1 s bins to facilitate visualization. **b**) GCaMP neuropil activation at 10-20 μm (red, solid line), 30-40 μm (orange, dashed line), 50-60 μm (green, –), 100-110 μm (cyan, –), 150-160 μm (blue, dotted line), and 245255 μm (violet, solid line) from the electrode site. Spatial and temporal neuropil falloff increase with frequency.

### Stimulation in somatosensory cortex activates neurons with different temporal activation behavior

Previous work evaluating the effects of microstimulation on neuronal activation utilized brief stimulation paradigms with pulse train durations on the order of milliseconds (Histed et al., 2009). However, during continuous stimulation, different temporal activation patterns were observed during the stimulation period in individual mice (Figs 3 & 4). Therefore, the aggregate temporal properties of neuronal activation across all mice (n=5) was examined. Temporal activation was separated into 1) onset neurons that rapidly decline in activity during stimulation and 2) steady-state neurons that remain activated throughout the stimulation period. Somas were classified as onset-responsive if they were activated during the first 0-2 s of the 30 s pulse train but ceased activation before the final 2 s of the pulse train (28-30 s); or as steady-state-responsive neurons if they were active during the final 2 s of stimulation (28-30 s post-stimulation onset) (Fig. 6a). In these acute experiments, all steady-state neurons were also active in the first 2 s of the pulse train. The duration of onset and steady-state neuron activity was measured by quantifying the total time during the stimulation period that GCaMP intensity was greater than an intensity threshold of three standard deviations from the pre-stimulus period. Further, steady-state neurons had a significantly greater Activation Time than onset neurons (Supplemental Fig. 2a, repeated measures two-way ANOVA with Bonferroni post-hoc comparison, F=374.831, p<0.001).

**Figure 6:**
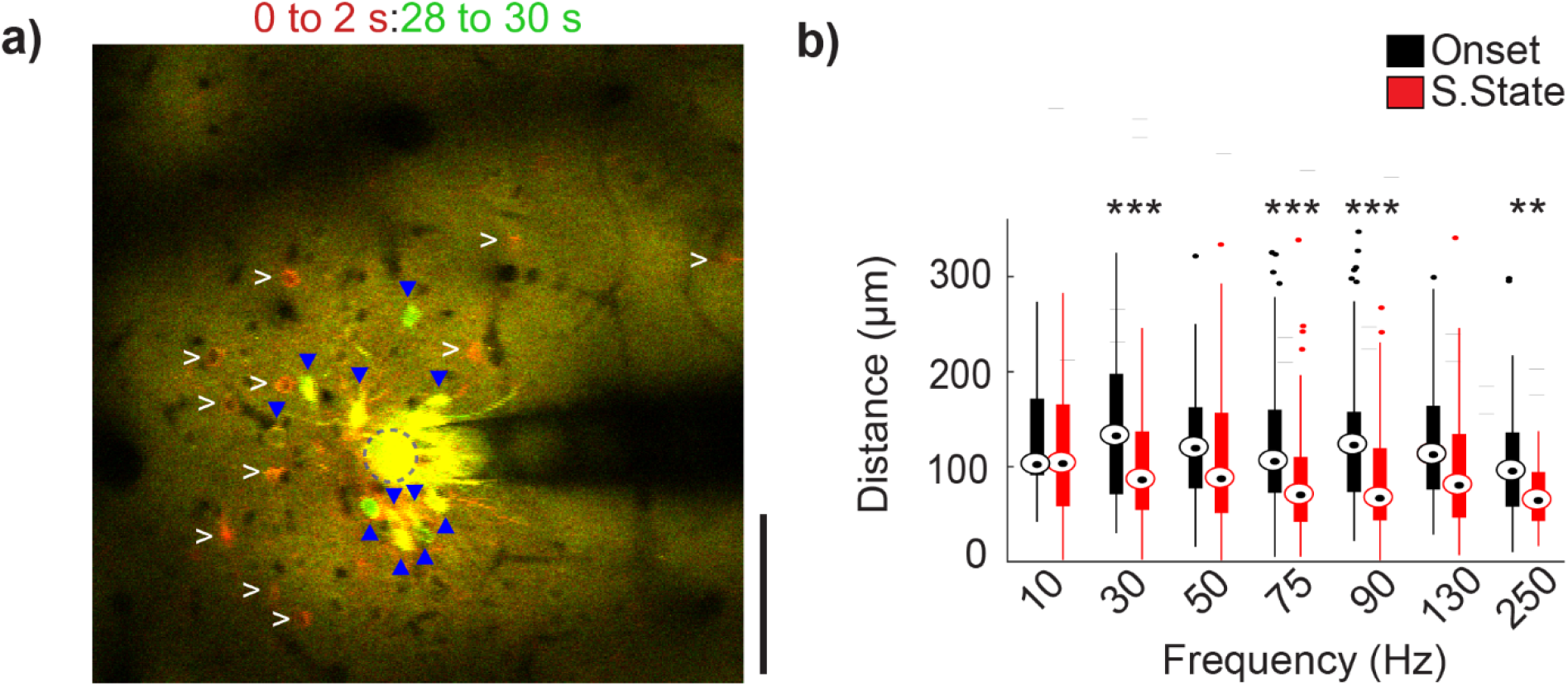
Onset and steady-state neurons are spatially distinct at high stimulation frequencies. (a) Mean GCaMP activity at the onset of the stimulation (0 to 2 s; red) and immediately before the end of stimulation (28 to 30 s; green) (stimulation parameter: 250 Hz, 50μA). Onset neurons (white arrowheads) and steady-state: (blue triangle) are marked in both images. Some steady-state neurons were also active at the onset of stimulation (yellow). (b) Average distance of the neurons activated during onset and steady-state phase. With increasing frequency, onset activated neurons are located significantly further from the electrode site than steady-state neurons. Repeated measures two-way ANOVA (within frequencies, n=15 neurons, F=14.707, p<0.005). Post-hoc t-tests with Bonferroni correction: **p<0.01 and ***p<0.001.

In order to evaluate if there was a spatial relationship for onset and steady-state somas, the spatial distribution of the two populations were compared (Fig. 6a). Since charge density accumulates on the edges of planar electrode sites, (Wei & Grill, 2009) distances were measured from the center of each soma to the nearest edge of the stimulating electrode site. Steady-state neurons were generally located closer to the electrode site than onset neurons, on average by ~40 μm (Fig. 6b). This phenomenon tended to occur at higher frequencies (Fig. 6b; at 130 Hz, p=0.058). Onset neurons and steady-state neurons showed distinct spatial relationships that were distal or proximal to the stimulation electrode, respectively.

### Stimulation in somatosensory cortex activates local and distant neurons in a frequency dependent manner

During continuous electrical stimulation at higher frequencies, onset neurons that decreased in activity tended to be at a greater distance from the electrode compared to steady state neurons (Fig. 5). We also evaluated the opposite relationship; if GCaMP activity in distant somas decreased over long stimulus durations at higher frequencies. While stimulation frequencies greater than 50 Hz led to onset and steady-state neurons exhibiting a confined spatial spread (Fig. 6b), not all onset neurons were distant neurons, and not all distant neurons were steady-state neurons. Therefore, in order to minimize bias, somas were also grouped by distance to evaluate if ‘local’ and ‘distant’ neurons exhibited distinct temporal relationships. The aggregated temporal activation patterns over local and distant neuronal populations in all mice (n=5) were considered. Somas were spatially classified by establishing a distance threshold (see methods, Fig. 7a), whereby cell bodies located within the threshold were considered ‘local’ neurons, while cells located outside the threshold were considered ‘distant’ neurons. No significant differences were observed between the number of activated local or distant neurons (Supplemental Fig. 3a).

**Figure 7:**
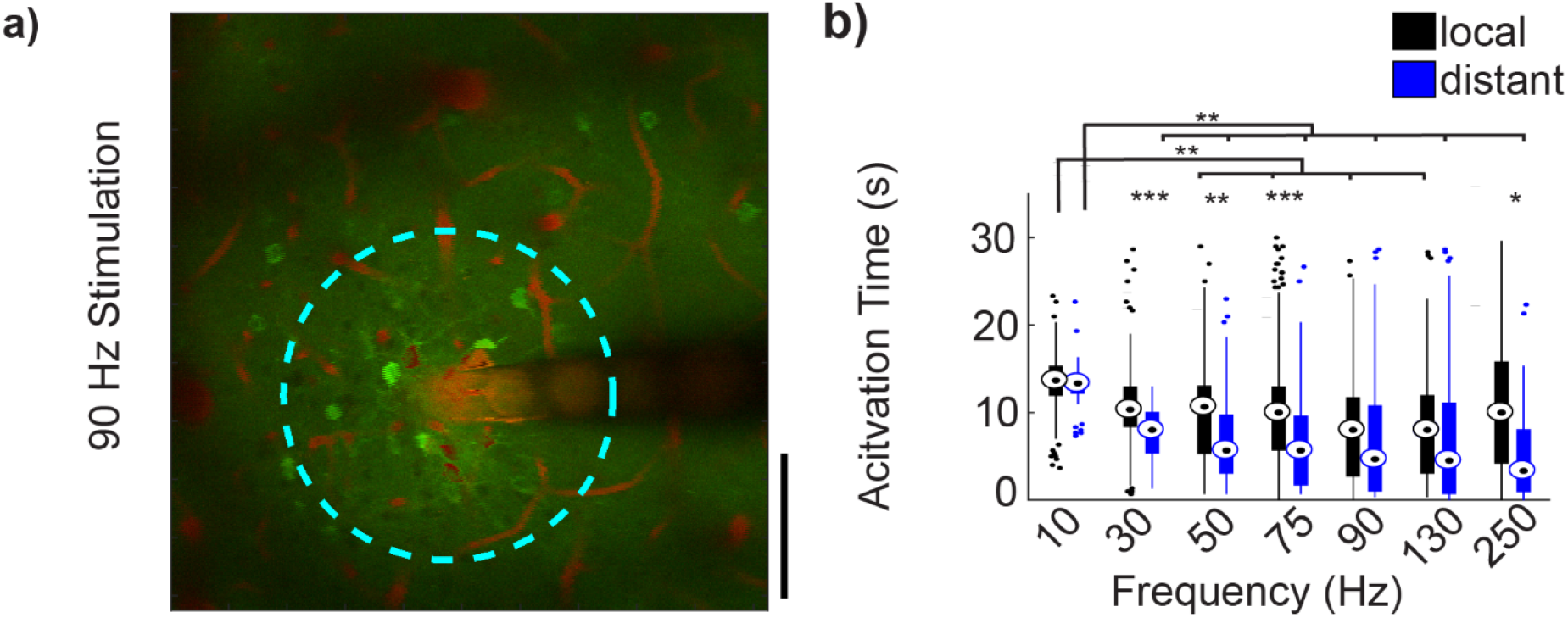
Activation properties of local and distant neuronal populations. (a) Standard deviation intensity projection over 30 seconds following 90Hz stimulation. Distance threshold shown by cyan circle. (b) Average activation time for local and distant activated neuron populations. Repeated measures two-way ANOVA (within frequencies, n=21 neurons, F=46.022, p<0.005; between frequencies F=10.483, p<0.005). Post-hoc t-tests with Bonferroni correction: *p<0.05 **p<0.01 and ***p<0.001.

Somatic calcium changes generally reflect action potential firing; although during periods of high spike rates, GCaMP6 fluorescence intensity can saturate due to the long decay constant (Chen et al., 2013). Because somas may exhibit initially high stimulation-evoked firing rates, which can be followed by a period of lower activity throughout the rest of the stimulation period (Fig. 5b; red), measurements of the duration of activation did not effectively capture the dynamics of neuronal responses. Therefore, to evaluate if local and distant somas had different temporal activation patterns, the ‘Activation Time’ of the two populations were compared. The Activation Time was defined as an estimate of the mean time at which the cell is responsive, weighted by the GCaMP intensity, and was calculated as the time-point at which the cumulative intensity reached 50% of the sum of the total ΔF/F_0_ signal change during the stimulation period. Activation Time was significantly greater in local somas than in distant somas for 30 Hz (10.9±0.4 vs 7.6±0.4 s), 50 Hz (9.8±0.6 vs 6.9±0.7 s), 75 Hz (10.5±0.6 vs 6.4±0.7 s), and 250 Hz (11.4±0.9 vs 5.7±1.5 s) stimulation (Fig. 7b, F=46.022, p<0.05). Additionally, Activation Time of both local and distant somas showed significant frequency dependence (Fig. 7b, F=10.482, p<0.001). Particularly of note was that for 10Hz stimulation, the local somas had significantly greater Activation Time than at higher stimulation frequencies (F=10.482, p<0.01).

As a spatial pattern of activation was observed during continuous stimulation (Fig 7b), the activation density for local and distant populations was measured. Activation density for local somas was calculated as the number of somas activated within the distance threshold, divided by the total area encompassed by the threshold. For distant somas, activation density was calculated using the area encompassed by an annulus with inner radius, r, equal to the distance threshold, and outer radius, *R*, equal to the distance of the furthest soma. For 10, 30, 75, and 90 Hz stimulation, local neurons had significantly greater neuronal density than distant somas (Supplemental Fig. 3b, F=2.993, p<0.05).

### Stimulation in somatosensory cortex creates a frequency dependent activation density pattern

In studies by Stoney et al, they noted activation of densely packed somas local to the electrode. In contrast, Histed et al noted only sparse activation of distant somas. To investigate this discrepancy, we evaluated the density of somas activated as a function of distance from the electrode. Prolonged electrical stimulation led to a dense activation pattern characterized by a decrease in activity in distant somas at frequencies >10 Hz. As before, somas were then separated into dense and sparse populations using a density threshold and evaluated for distinct spatial and temporal relationships. In order to evaluate if continuous electrical stimulation can activate a set of cells restricted to a small spatial area, patterns of somatic activation were classified as dense or sparse. Concentric circular bins were drawn with their origin at the center of the electrode site with subsequent bins increasing in 20 μm radial increments (Fig. 8a). Density was calculated as the number of activated neurons in each bin, divided by the concentric bin area (Fig 8b). Density thresholds were established similarly to distance thresholds.

**Figure 8:**
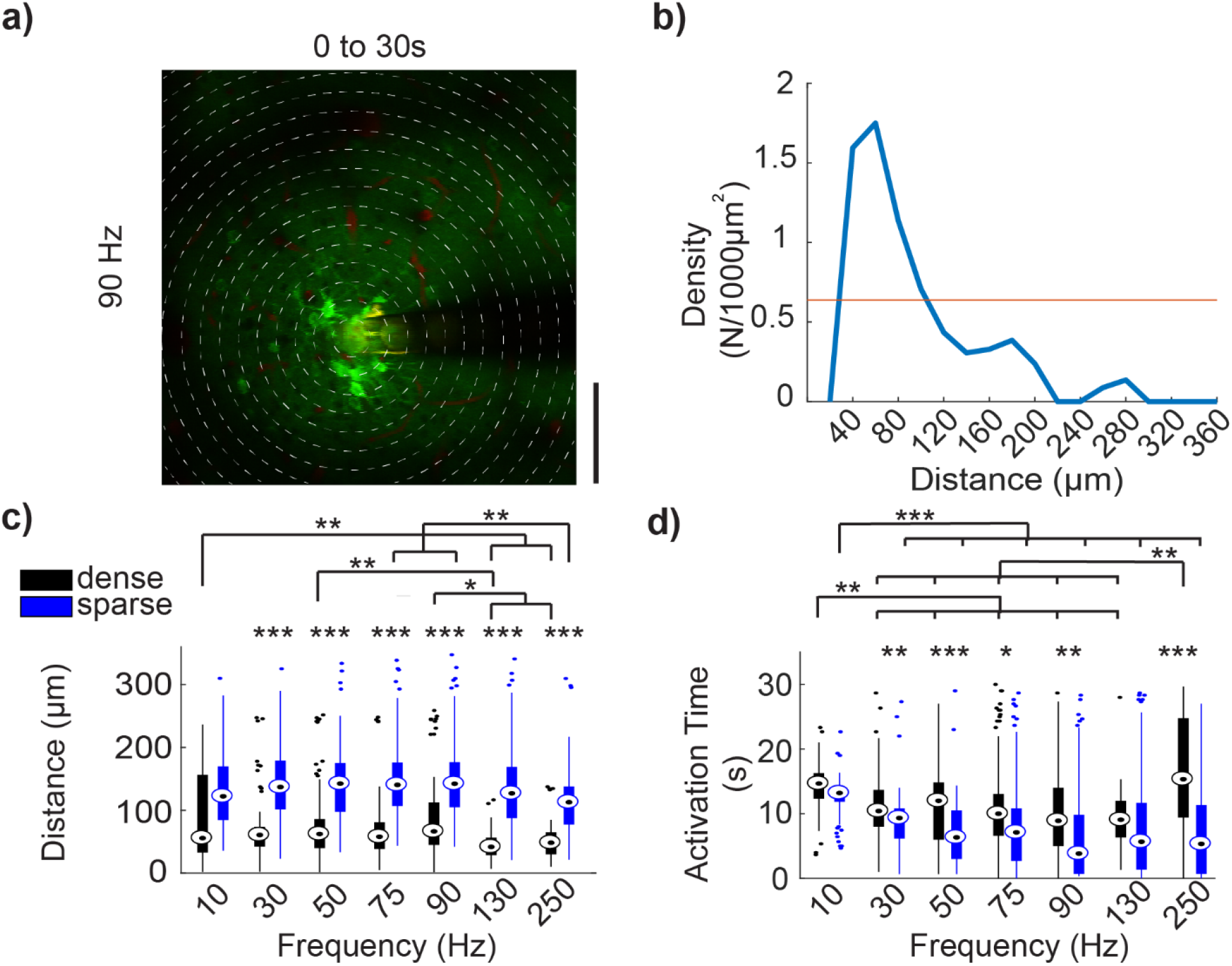
Activation properties of dense and sparse populations of neurons. (a) Mean GCaMP activity (30 s, 90 Hz stimulation) with concentric bins of 20 μm incremental radius around the center of the electrode site, demonstrating density calculation method. (b) For the same trial, density (N/1000 μm^2^) is plotted against distance of the bins from the electrode. (c) Average distance of dense and sparse activated neurons. For all stimulation frequencies, sparsely activated neurons are located farther from the electrode site than dense activated neurons. (d) Average activation time of dense and sparse activated neuron population. Dense activated neurons are active for a longer duration in all stimulation frequency except 130Hz. Repeated measures two-way ANOVA (for (c) within frequencies: n=42 neurons, F=180.332, p<0.001; between frequencies: F=28.137, p<0.001. for (d) within frequencies: n=42 neurons, F=15.088, p<0.001; between frequencies: F=5.527, p<0.001). Post-hoc t-tests with Bonferroni correction: *p<0.05, **p<0.01 and ***p<0.001

Activated somas within dense populations were located significantly closer to the electrode site for all stimulation frequencies except 10 Hz (Fig. 8c, F=180.332, p<0.001). Additionally, densely activated somas generally showed significantly greater Activation Times compared to sparsely activated somas (Fig. 8d, F=15.088, p<0.05). The Activation Time of dense and sparse somas showed similar frequency dependence as observed in local and distal somas in Fig. 7b. However, the Activation Time for dense neurons at 250 Hz stimulation was significantly greater than at stimulation frequencies tested from 30-130 Hz (F=5.527, p<0.01). Somas that were classified as higher density were closer to the electrode and had longer Activation Times compared to sparse cells, which were further from the electrode and had shorter Activation Times. At the onset of electrical stimulation, there was a low density, sparse, and distributed activation pattern of neurons.

## Discussion

This work shows that continuous electrical stimulation for 30 seconds in acute experiments strongly activated somas located in close proximity to the electrode for the duration of the stimulation period for frequencies >10 Hz (Figures 2–8). Previous work by Histed et al utilizing 2-photon microscopy and a calcium indicator demonstrated that stimulation with <1 s pulse trains of cathodic phase leading, 200 μs/phase, biphasic pulses under 10 μA in current at 250 Hz, resulted in a sparse, distributed population of activated neurons surrounding the electrode tip, without a strong bias towards neurons near the tip (Histed et al., 2009). Increasing current over the 4-9 μA range increased the density of neurons activated within this 100-200 μm sphere, again without a bias towards closer to the stimulation site (Histed et al., 2009). Through multiple experimental manipulations they concluded that stimulation in this confined parameter space only directly activates axons within 30 μm of the electrode, which in turn antidromically activates the somas associated with these axons, yielding the observed sparse pattern of somatic activation (Histed et al., 2009). These results were counter to prevailing theory based on seminal studies by Stoney et al which suggest that stimulation leads to a sphere of activated neurons near the electrode tip, with the size of the sphere growing in proportion to the current applied (Murasugi, Salzman, & Newsome, 1993; Ranck, 1975; Rattay, 1999; Robinson & Fuchs, 1969; Stoney Jr et al., 1968; Tehovnik, 1996; Tolias et al., 2005).

The Histed results were obtained using stimulation charges at or just above the threshold for eliciting a significant response using a calcium indicator (0.8 to 1.8 nC/phase), applied for brief periods (<1 s) and at very high frequency (250 Hz), which led to the implication that, “it is impossible to activate a set of cells restricted to a small spatial volume” (Histed et al., 2009). In contrast, the results described in our study using longer pulse trains (30 s) and larger stimulation charges (2.5 nC/phase) show a distinct pattern of somatic activation across frequencies consisting of a comparatively denser group of somas near the electrode tip and a sparse group of neurons located distant from the electrode. In short, our results support both the original studies by Stoney and the more recent study by Histed, painting a complex picture of spatial activation that is dependent on charge applied and time of observation during an extended pulse train (Histed et al., 2009; Stoney Jr et al., 1968).

The data in the present study demonstrate that, in general, activated somas both local and distant remain activated at lower stimulation frequencies throughout the duration of the pulse train. At higher frequencies (90 Hz and above), distant somas activated at the beginning of the 30 s pulse train cease to remain active by the end of the pulse train, with the Activation Time progressively shorter as the frequency was increased. The responses of local axons versus somas also differentially changed over the course of an extended pulse train, where frequencies over 50 Hz lead to initial transient activation that eventually subsides over the course of the pulse train.

The idea that different elements of the neuron may be activated or inactivated at different applied currents is not new (Cameron C McIntyre & Grill, 2002; Overstreet, Klein, & Tillery, 2013). It is often assumed, however, that at frequencies at which pulse to pulse entrainment is feasible (130 Hz or less), the differential evoked responses local to the electrode are primarily a function of charge applied (current*pulsewidth), given that stimulation is within the Shannon/McCreery safety limits (Cogan et al., 2016; Shannon, 1992). In this paradigm, changes in evoked network response as a function of frequency and time over the duration of pulse train have been presumed to be presynaptically mediated, e.g. vesicular depletion at the synapse leading to synaptic depression (Y. Wang & Manis, 2008). However, stimulation of CA1 neurons via antidromic activation of efferent fibers exhibits time-varying responses, where the transient population response at the onset of stimulation exceeds steady state activity during stimulation (Cai et al., 2017). Notably, the magnitude of the antidromically evoked population responses was found to be frequency dependent, where higher frequencies of stimulation elicited weaker onset and steady state responses. Moreover, high frequency stimulation of axons at 100 and 200 Hz disrupts the synchronicity of evoked neuronal responses compared to 50 Hz (Z. Wang, Feng, & Wei, 2018). Very high frequency stimulation (~10 kHz) can block neural activity altogether in isolated fibers in the periphery. While McCreery et al. has demonstrated that continuous stimulation for periods of only seven hours can lead to Stimulation Induced Depression of Neuronal Excitability (SIDNE) (D. B. McCreery, Agnew, & Bullara, 2002; D. B. McCreery, Yuen, Agnew, & Bullara, 1997), and other studies have shown that continuous stimulation for as short as 1 hr/day could lead to significantly different behavioral outcomes (Hao et al., 2015; White & Sillitoe, 2017), the activation and then subsequent decrease of activation in local axons and their connected soma over the course of a much shorter 30 second pulse train, suggests an underlying mechanism that has not been previously explored.

The mechanism by which increasing frequency may cause transient activation of axons local to the electrode followed by inactivation over the pulse train is presently not understood. Continuous stimulation could cause intermittent failure to propagate action potentials through the axon at frequencies as low as 30 Hz, putatively mediated by extracellular accumulation of K+ ions, depletion of ATP, and reduced inward Na+ current (Smith, 1980). Alternatively, it is possible that increasing frequency could lead to virtual anode generation. Increasing intensities of cathodic stimulation may generate a virtual anode which hyperpolarizes membranes on stimulated fibers distant from the cathode, and blocks the propagation of action potentials initiated near the cathode (Ranck, 1975). Due to the capacitive nature of the electrode-electrolyte interface and the lipid bilayer of the neural element (Bates & Chu, 1992), increasing stimulation frequency at a fixed amplitude will yield a lower impedance that may facilitate virtual anode generation. Further studies are necessary to better understand the cause of inhibition of calcium activity in distal somas at higher frequencies.

Under normal conditions, neuronal depolarization causes an influx of intracellular calcium through pathways such as voltage or neurotransmitter gated Ca^2^+ channels, which is measured by the GCaMP indicator (Berridge, 1998). The calcium influx then leads to Ca^2^+ release through channels present in intracellular stores, mainly the endoplasmic reticulum (Sharp et al., 1993). Intracellular calcium can then induce local changes at the synaptic level or regulate gene expression (Berridge, 1998). However, excessive levels of intracellular calcium can have neurotoxic effects (Arundine & Tymianski, 2003). We observed activated cells near the stimulating electrode site with sustained GCaMP fluorescence without having discernable spikes (as observed in the plateau in fluorescence change in Fig. 5). This may be due to the simultaneous activation of inhibitory neurons (non-GCaMP expressing) (Dana et al., 2014). Nevertheless, the long-term impact that sustained intracellular calcium concentrations induced by microstimulation may have on local gene expression, synaptic properties, and neuronal health remains to be elucidated. For example, continuous stimulation at parameters above the safety limits (k>1.7~1.85) established by the Shannon curve can cause damage to tissue in the vicinity of the electrode, which is often attributed to driving toxic electrochemical reactions at the surface of the electrode, or driving local neural tissue to increase activity beyond physiological norms leading to excitotoxic effects (Agnew, Yuen, Pudenz, & Bullara, 1975; Cogan et al., 2016; Merrill, Bikson, & Jefferys, 2005; Mortimer, Shealy, & Wheeler, 1970; Shannon, 1992; Yuen, Agnew, Bullara, Jacques, & McCreery, 1981). Furthermore, pioneering work by McCreery suggests that 4 nC/phase represents a stimulation threshold above which obvious damage occurs, as assessed by gross histological metrics, as opposed to larger electrodes which are governed by the Shannon equation (D. McCreery, Pikov, & Troyk, 2010; Douglas B McCreery, Agnew, Yuen, & Bullara, 1990). While this present study was carried out below the Shannon limit and charge density per phase outlined by McCreery, the biological substrates of plasticity induced by clinically relevant stimulation parameters over time remain poorly understood.

Additionally, various studies have demonstrated that electrical stimulation can cause diverse effects such as increases in hippocampal and thalamic volume, increases in blood vessel size and synaptic density, cortical dendrite growth, and changes in mRNA expression - some of which may influence short term local somatic and axonal response (Agnew et al., 1975; Baudry, Oliver, Creager, Wieraszko, & Lynch, 1980; Brownson, Little, Jarvis, & Salmons, 1992; Chakravarty et al., 2016; Cooperrider et al., 2014; Gall, Murray, & Isackson, 1991; Hirano, Becker, & Zimmerman, 1970; Morris, Feasey, ten Bruggencate, Herz, & Hollt, 1988; Pudenz, Bullara, Dru, & Talalla, 1975; Sankar et al., 2015; Sankar et al., 2016; Veerakumar et al., 2014; Xia, Buja, Scarpulla, & McMillin, 1997; Xia et al., 2000). In chronic neuromodulation or neuroprosthetic applications, additional complexities arise from tissue changes in both the neuronal and non-neuronal populations due to the implantation injury (glia, neurovasculature, immune cells) (Kolarcik et al., 2015; Takashi Daniel Yoshida Kozai, Jaquins-Gerstl, Vazquez, Michael, & Cui, 2015; Michelson et al., 2018; Salatino, Ludwig, Kozai, & Purcell, 2017; Steven M. Wellman & Kozai, 2017), electrode material degradation (Alba, Du, Catt, Kozai, & Cui, 2015; Cogan, 2008; Cogan et al., 2016; T. Kozai et al., 2014; Steven M Wellman et al., 2018; Wilks et al., 2017), and plastic changes from long-term electrical stimulation (Agnew et al., 1975; Baudry et al., 1980; Brownson et al., 1992; Chakravarty et al., 2016; Cooperrider et al., 2014; Gall et al., 1991; Hirano et al., 1970; Merrill et al., 2005; Morris et al., 1988; Pudenz et al., 1975; Sankar et al., 2015; Sankar et al., 2016; Veerakumar et al., 2014; Xia et al., 1997; Xia et al., 2000). Future studies should be aimed at disentangling the complex network of dendrites, axons, and cell bodies of inhibitory and excitatory cells that may be present near the electrode in the brain (Overstreet et al., 2013).

The frequency dependent inactivation of axons local to the electrode over the course of an extended pulse train may provide insight on a potential mechanism of action for electrical stimulation for specific therapeutic uses (Agnesi, Muralidharan, Baker, Vitek, & Johnson, 2015; Gildenberg, 2005). For example, as DBS of the subthalamic nucleus (STN) to treat the tremor associated with Parkinson’s Disease was originally based on the positive results of lesioning the same area, STN DBS was originally thought to generate the equivalent of a titratable/reversible lesion of STN. Following studies utilizing microwires for electrophysiological recordings – which are biased towards sampling action potentials initiated at the axon hillock, as this is where the largest current deflection occurs – suggested that therapeutic DBS parameters in the 130 Hz range differentially suppress the activity of local STN somas while still activating local axons projecting to the targets from STN nuclei (Cameron C. McIntyre, Grill, Sherman, & Thakor, 2004). This led to the suggestion that DBS may be creating an information lesion of signals passing through STN, as entrainment at 130 Hz would prevent the passage of physiological signals with any temporal information, due to the stimulus rate and the absolute and relative refractory period of a neuron (Agnesi, Connolly, Baker, Vitek, & Johnson, 2013; Grill, Snyder, & Miocinovic, 2004). More recently antidromic activation of tracks of passage traveling near STN from motor cortex, known as the hyperdirect pathway, have received increasing attention as a potential mechanism to ameliorate pathological thalamocortical oscillations driving tremor in Parkinson’s Disease (Gradinaru, Mogri, Thompson, Henderson, & Deisseroth, 2009; Li, Arbuthnott, Jutras, Goldberg, & Jaeger, 2007; Cameron C McIntyre & Hahn, 2010). In contrast, 10 Hz optogenetic stimulation in the brain leads to the highest probability of entrainment, with the entrainment probability significantly decreasing at frequencies greater than 15 Hz (Bitzenhofer et al., 2017). Therefore, it may not be surprising that 10 Hz electrical stimulation leads to continuous GCaMP activity over long stimulus pulse trains. The results in the present study demonstrate that stimulation at high frequencies associated with therapeutic DBS initially activates both local soma and local axons connected to distant somas, but over time the evoked activity in local axons projecting to distant somas decreases. These data suggest that to understand the post-synaptic response to stimulation – and ultimately the therapeutic mechanisms of action – one must account for differential activation of axons vs somas near the electrode, as well as changes in the local activation profile as a function of frequency and time.

### Limitations

Although care was taken to allow neuronal fluorescence intensities to return to pre-stimulation levels before another frequency was tested, stimulation frequencies were evaluated in increasing frequencies instead of randomizing the frequency order. Future studies should randomize one stimulation frequency to each implant location as well as evaluate the impact of the total number of stimulation pulses compared to stimulation frequency on driving stimulation induced depression of neuronal excitability. However, in two animals, 10 Hz trials were repeated after the entire frequency sequence and remained consistent with the first 10 Hz trial. Another key limitation is that calcium activity was observed under ketamine anesthesia, which inhibits NMDA receptors and potentiates GABA_A_ receptors (Alkire, Hudetz, & Tononi, 2008), and shifts the balance of excitatory and inhibitory transmission in cortex (Haider, Hausser, & Carandini, 2013; Homayoun & Moghaddam, 2007; Michelson & Kozai, 2018). These changes in neuronal excitability and network activity might potentially impact the frequency dependent responses to microstimulation observed here. Finally, all experiments were carried out on male mice in this limited scope study. Future work will include comparisons to female mice.

## Conclusion

Despite the widespread use of electrical stimulation as a scientific tool and a modality to affect therapeutic outcomes, a lack in understanding of its effects on network-level neural activity has limited its applicability. Without complete understanding of how an exogenous electric field is interacting with neuronal and nonneuronal cells local to a chronically implanted electrode at the targeted point of intervention – and activating or inhibiting individual elements of these cells types – it is difficult to understand the mechanism by which targeted electrical stimulation may be creating desired therapeutic outcomes or undesirable side effects. While the present work is not a complete study on the large parameter space of electrical stimulation, we disprove the hypothesis that the sparse and distributed neural activation patterns occur under continuous stimulation in a frequency independent manner. Instead, different stimulation parameters can dramatically alter the activation pattern at the cellular level in cortex. A deeper understanding of electrical stimulation parameters on neuronal activation (and non-neuronal activation) will be critical in utilizing electrical stimulation for targeted applications in basic science research as well as clinical therapeutics. Thus, the current work motivates additional non-clinical, non-translational basic science research to elucidate the mechanisms governing electrical stimulation.

## Acknowledgements

We thank Riazul Islam for assistance with neuron ROI selection, and James Trevathan and Evan Nicolai from the Mayo Clinic for helpful discussions. This work was supported by NIH NINDS R01NS094396, R01NS094404, R01NS062019, R01NS089688, and R21NS108098.

## Author Contributions

All authors had full access to all the data in the study and take responsibility for the integrity of the data and the accuracy of the data analysis. *Conceptualization*, T.D.Y.K., A.L.V., and K.A.L.; *Methodology*, N.J.M. and T.D.Y.K.; *Investigation*, J.R.E. and T.D.Y.K.; *Formal Analysis*, N.J.M. and J.R.E.; *Writing – Original Draft*, T.D.Y.K., K.A.L., and N.J.M.; *Writing – Review & Editing*, all authors; *Visualization*, N.J.M., and T.D.Y.K.; *Supervision*, T.D.Y.K.; *Funding Acquisition*, T.D.Y.K.

## Supplemental File

**Supplemental Figure 1.**
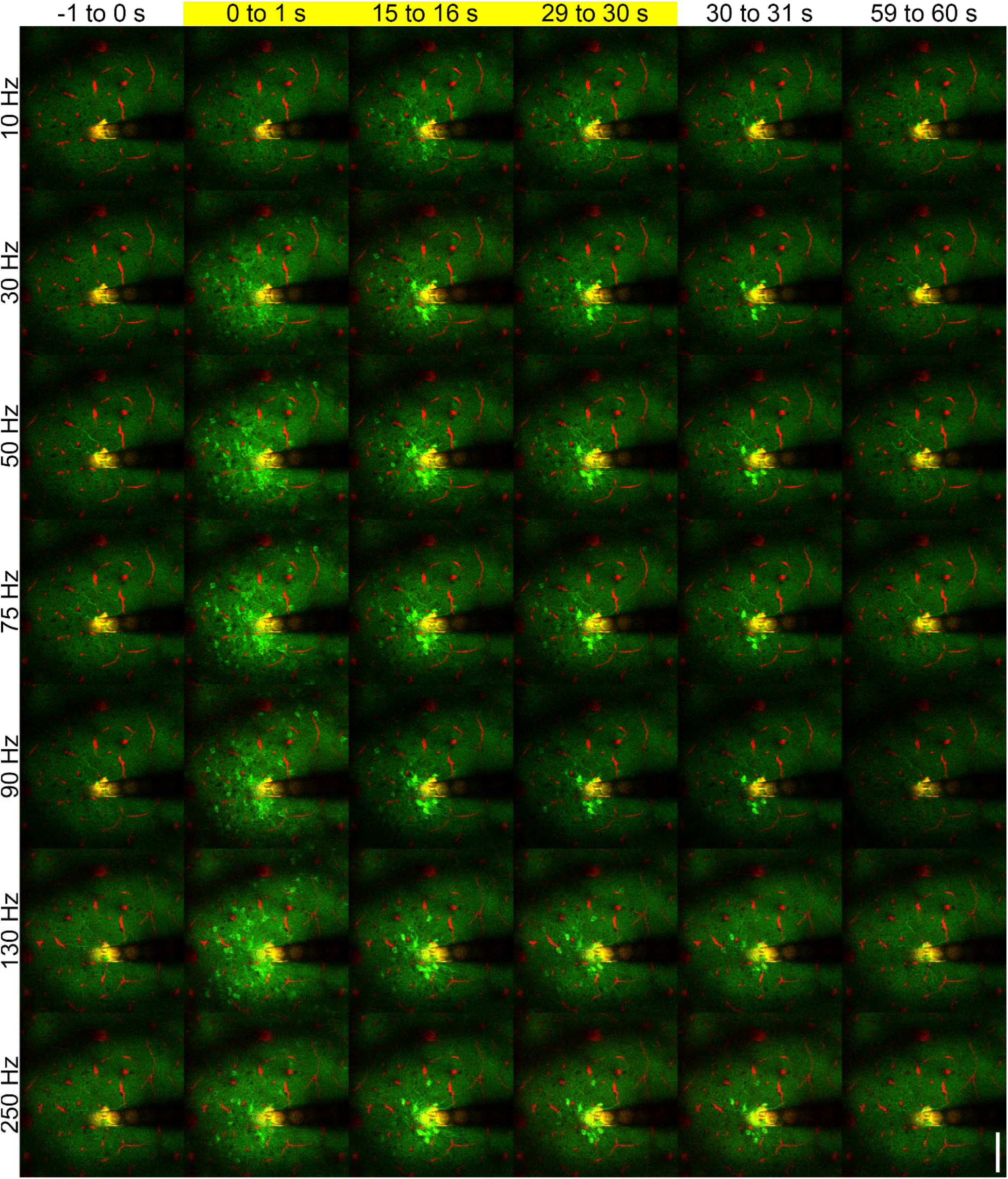
Same as figure 3 for all frequencies. Scalebar = 100 μm

**Supplemental Figure 2.**
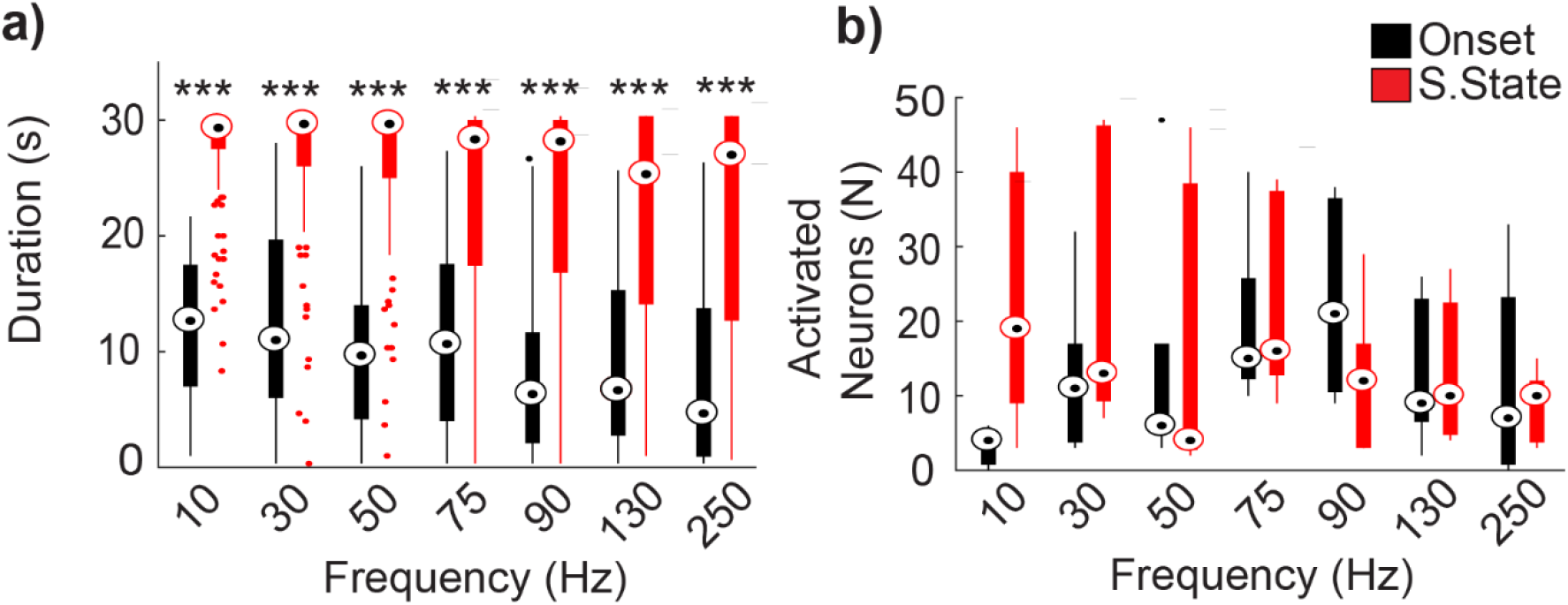
(a) Average duration of onset and steady-state phase. Duration of activation phases were calculated by tracking onset-responsive (neurons that are active 0-2s into stimulation) and steady-state-responsive neurons (neurons active 28-30s into stimulation). (b) Average number of neurons activated during onset and steady-state phase for 10Hz, 30Hz, 50Hz, 75Hz, 90Hz, 130Hz and 250 Hz stimulation. For (a): repeated measures two-way (within frequencies, F=374.831, p<0.001) with post-hoc t-tests with Bonferroni correction ***p < 0.001.

**Supplemental Figure 3.**
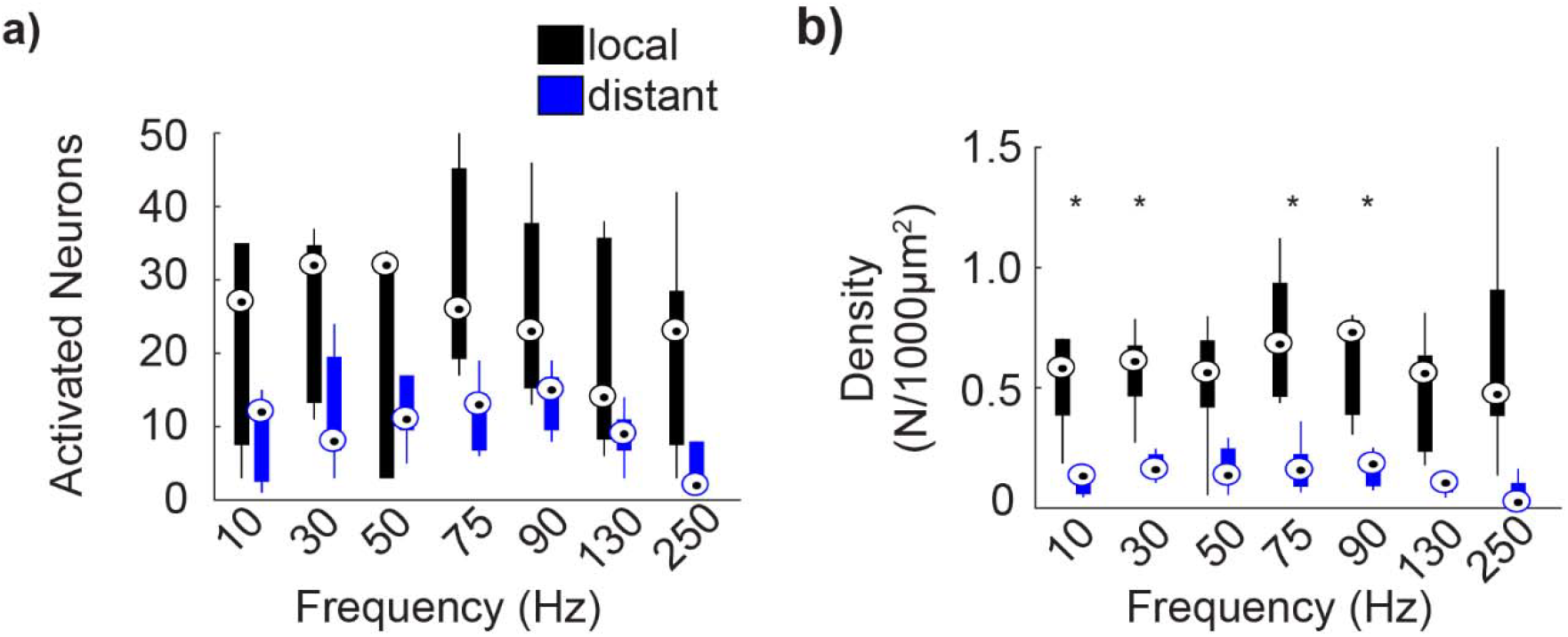
(a) Average number of neurons activated that are local and distal from the electrode. (b) Average activation density of local and distal neuron population. Activation density for locally activated neurons was significantly higher for all stimulation frequencies than distally activated neurons. For (b): repeated measures two-way ANOVA (within frequencies, F=523.846, p<0.001) and post-hoc t-tests with Bonferroni correction *p < 0.05

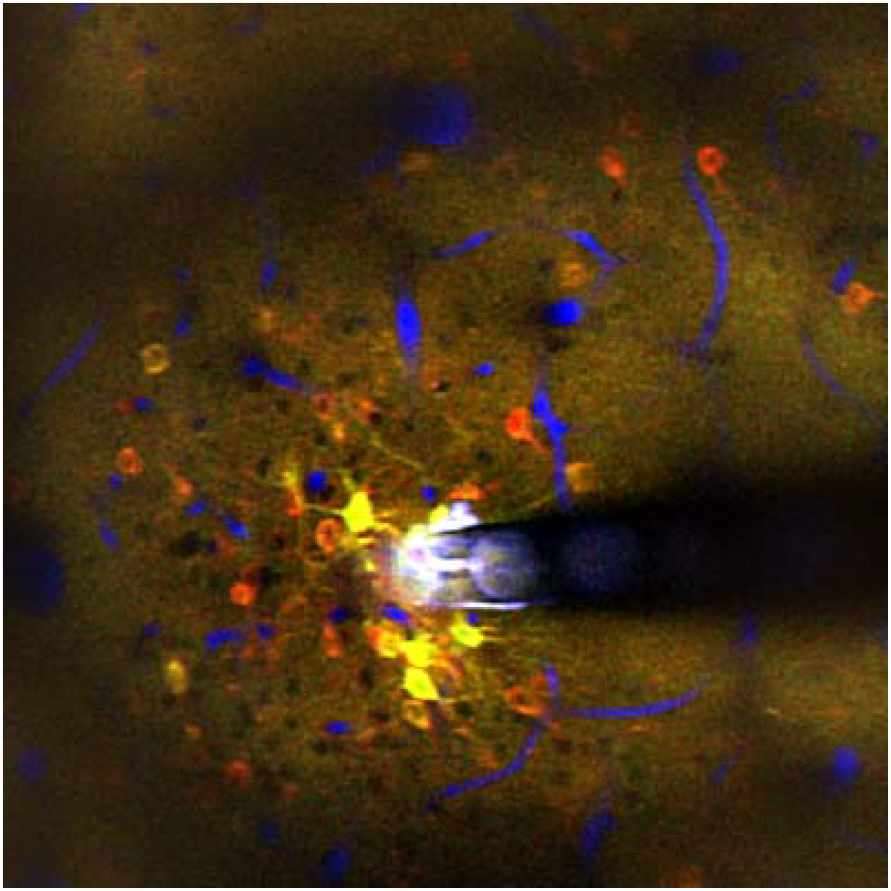

Neuronal microstimulation can treat neurological disorders, but the therapeutic mechanism is not well understood. Microstimulation in mouse cortex produced a spatiotemporal response, activating a sparse population of neurons only at the onset of stimulation (red), and a dense population of neurons throughout the stimulation duration (yellow). Vessels shown in blue.

